# Anticancer effect of zanubrutinib in HER2-positive breast cancer cell lines

**DOI:** 10.1101/2023.02.23.529815

**Authors:** Hana Dostálová, Radek Jorda, Eva Řezníčková, Vladimír Kryštof

**Affiliations:** Department of Experimental Biology, Faculty of Science, Palacký University Olomouc, Šlechtitelů 27, 78371 Olomouc, Czech Republic; Institute of Molecular and Translational Medicine, Faculty of Medicine and Dentistry, Palacký University Olomouc, Hněvotínská 5, 77900 Olomouc, Czech Republic

**Keywords:** BTK inhibitor, drug repurposing, HER2 positive breast cancer, zanubrutinib

## Abstract

Small molecule Bruton’s tyrosine kinase (BTK) inhibitors have been developed for the treatment of various haemato-oncological diseases, and ibrutinib was approved as the first BTK inhibitors for anticancer therapy in 2013. Previous reports proved the receptor kinase human epidermal growth factor receptor 2 (HER2) to be a valid off-target kinase of ibrutinib and potentially other irreversible BTK inhibitors, as it possesses a druggable cysteine residue in the active site of the enzyme. These findings suggest ibrutinib as a candidate drug for repositioning in HER2-positive breast cancer (BCa). This subtype of breast cancer belongs to one of the most common classes of breast tumours, and its prognosis is characterized by a high rate of recurrence and tumour invasiveness. Based on their similar kinase selectivity profiles, we investigated the anticancer effect of zanubrutinib, evobrutinib, tirabrutinib and acalabrutinib in different BCa cell lines and sought to determine whether it is linked with targeting the epidermal growth factor receptor family (ERBB) pathway. We found that zanubrutinib is a potential inhibitor of the HER2 signalling pathway, displaying an antiproliferative effect in HER2-positive BCa cell lines. Zanubrutinib effectively inhibits the phosphorylation of proteins in the ERBB signalling cascade, including the downstream kinases Akt and ERK, which mediate key signals ensuring the survival and proliferation of cancer cells. We thus propose zanubrutinib as another suitable candidate for repurposing in HER2-amplified solid tumours.

## Introduction

HER2-positive breast cancer is one of the most frequent types of tumours in women, accounting for nearly 20–30% of breast cancer cases. The amplification or overexpression of the *HER2* gene as well as its amenability to pharmacological modulation makes it an attractive target for drug discovery. The HER2 kinase is a member of the ERBB tyrosine kinase receptor family, together with EGFR, HER3 and HER4. Despite their molecular homology, ERBB kinases affect different signalling pathways depending on their homo- and heterodimerization, which is crucial for their enzymatic activity [1]. The HER2 pathway contributes to the regulation of cell growth, proliferation and survival via various signalling cascades, including the Ras/Raf/MAPK, mTOR and PI3K/Akt pathways [2]. Oncogenic activation of HER2 contributes to deregulated proliferation of breast tissue cells, leading to tumorigenesis. Among the molecular subtypes, HER2-positive breast cancer has a poor prognosis, with a higher rate of recurrence and tumour invasiveness [3].

However, HER2-positive carcinomas benefit from standard therapy combined with targeted therapy using anti-HER2 antibodies and small molecule inhibitors. The first approved monoclonal antibody, trastuzumab, acts through multiple mechanisms. Upon binding of the drug to the receptor, the HER2 downstream RAS/MAPK and PI3K/AKT signalling pathways are inhibited, and ubiquitin-mediated degradation of HER2 is induced [4]. Another possible mechanism of action is through the attraction of innate immune cells [5]. Another example of an anti-HER2 agent is pertuzumab, which is frequently used in targeted therapy together with trastuzumab. The combination of these two antibodies appears to show high efficacy, as the inhibitory mechanism of pertuzumab complements that of trastuzumab by binding to a different epitope in the extracellular domain of the HER2 receptor [6]. Although acquired resistance to targeted therapies is a common obstacle in setting up a treatment regimen, insensitivity can be overcome by using small molecule inhibitors such as lapatinib, neratinib or tucatinib that target the catalytic function of the kinase [7, 8]. In addition, new drugs are still being sought. Apart from traditional drug discovery approaches, alternative ways include drug repurposing, i.e., searching for drugs among already approved pharmaceuticals.

Bruton’s tyrosine kinase (BTK) inhibitors are a group of low-molecular-weight kinase inhibitors that are effective in certain B-cell malignancies. Their target, the nonreceptor kinase BTK, is a key effector in the B-cell receptor pathway. Ibrutinib is a first-in-class drug that has been approved for the treatment of chronic lymphocytic leukaemia and the non-Hodgkin’s lymphomas follicular lymphoma, mantle cell lymphoma, and Waldenström’s macroglobulinemia [9]. Ibrutinib is a covalent inhibitor that reacts with Cys481 in the active site of the kinase. A variety of second-generation BTK inhibitors with improved pharmacological properties have been developed, including acalabrutinib, zanubrutinib, tirabrutinib and evobrutinib [9]. Their biochemical properties were compared in a review article by Estupiñán et al., 2021 [10].

In addition to their primary use in B-cell malignancies, repurposing of BTK inhibitors beyond the primary indications is being extensively investigated [11–13]. The drugs could be used for therapy of other cancer types expressing BTK, and some of them are even currently being investigated in clinical trials (e.g., in acute myeloid leukaemia [14], colorectal carcinoma [15], prostate cancer [16]). In addition, the off-target kinases of BTK inhibitors provide a possibility to expand the use of these inhibitors in other cancers. Some of these kinases belong to two major families, namely, TEC (ITK, TEC, BMX and RLK/TXK) and ERBB (EGFR, HER2 and HER4) [17]. These kinases share a suitably positioned cysteine residue (analogous to Cys481 in BTK) in the active site. Importantly, ibrutinib and acalabrutinib have been identified to potently block ERBB and TEC kinases expressed in solid tumours [18–20]. Currently, there are over 20 ongoing clinical trials to verify the therapeutic efficacy of these two drugs beyond haemato-oncological malignancies, either in combination therapies or, more interestingly, as monotherapies.

The BTK inhibitor zanubrutinib has been approved for the treatment of mantle cell lymphoma and Waldenström’s macroglobulinemia [21]. Regarding its use in haemato-oncological malignancies, zanubrutinib has similar or slightly improved selectivity towards kinases in the TEC and ERBB families compared to ibrutinib, yet it is less potent than acalabrutinib [10]. However, its kinase selectivity profile has revealed promising potency towards ERBB kinases, namely, EGFR and HER4 [22, 23] (Supplementary Table 1). In fact, its biochemical properties are similar to those of ibrutinib [9, 10, 22], indicating that it could be another suitable candidate for repurposing in HER2-amplified solid tumours. Among the next-generation BTK inhibitors, zanubrutinib possesses the lowest IC50 values for the HER2 receptor [10]. We therefore investigated the anticancer effect of zanubrutinib in breast cancer cell lines. We describe zanubrutinib as a potential inhibitor of the HER2 signalling pathway, displaying antiproliferative effects in HER2-positive breast cancer cell lines, and we propose zanubrutinib as a candidate drug to be further investigated as a therapeutic agent in HER2-amplified breast cancer.

## Methods

### Cell lines and compounds

Human cancer cell lines (obtained from ATCC, USA, or DSMZ, Germany) were cultured according to the distributors’ instructions. In brief, MCF7, SKBR3, BT20, JIMT1 and BT474 cells were maintained in Dulbecco’s modified Eagle’s medium supplemented with 10-15% FBS, and T47D, HCC1806 and EFM192A cells were maintained in RPMI 1640 medium supplemented with 10-20% FBS. All media were supplemented with 100 U/mL penicillin, 100 μg/mL streptomycin, and 2 mM glutamine. For treatment, cells were seeded at densities of 1.5-2 million cells per dish in 60 mm dishes and allowed to adhere overnight. The BTK inhibitors ibrutinib, evobrutinib, tirabrutinib, acalabrutinib and zanubrutinib were purchased from MedChemExpress, and the EGFR/HER2 inhibitor lapatinib was purchased from LC Laboratories.

### Cytotoxicity assay

For the cytotoxicity assays, cells were seeded into 96-well plates. After overnight preincubation, cells were treated in triplicate with six different concentrations of each compound for 72 h. After treatment, resazurin solution (Sigma Aldrich) was added for 4 h, and the fluorescence of resorufin, corresponding to living cells, was measured at 544 nm/590 nm (excitation/emission) using a Fluoroskan Ascent microplate reader (Labsystems. The results of the assays were used to construct dose-response curves, and GI50 values (the drug concentration lethal to 50% of the cells) were calculated.

### Immunoblotting

After cell lysis, proteins were separated using SDS-PAGE and electroblotted onto a nitrocellulose membrane. After 1 hour of blocking with bovine serum albumin, the membrane was incubated overnight with specific primary antibodies and then for 1 hour with peroxidase-conjugated secondary antibodies. Peroxidase activity was then detected with SuperSignal West Pico reagents, and band intensities were measured using a LAS-4000 CCD camera. The following specific antibodies were used and purchased from Cell Signaling: anti-EGFR (D38B1), anti-HER2/ErbB2 (D8F12), anti-HER3/ErbB3 (D22C5), anti-HER4/ErbB4 (111B2), anti-phospho-EGFR Y1068 (D7A5), anti-phospho-HER2/ErbB2 Y1221/1222 (6B12), anti-phospho-HER3/ErbB3 Y1289 (D1B5), anti-phospho-HER4/ErbB4 Y1284/EGFR Y1173 (21A9), anti-Akt (pan) (C67E7), anti-phospho-Akt S473 (D9E), anti-p44/42 MAPK (Erk1/2), anti-phospho-p44/42 MAPK (phospho-Erk1/2) T202/Y204), and anti-PARP (46D11). Anti-β-actin (C4) was purchased from Santa Cruz Biotechnology.

### Cell cycle analysis

Analysis of the cell cycle distribution was performed in 96-well plates. Asynchronous cells were seeded and treated with different concentrations of the compounds for 24 hours. After incubation, 5x staining solution supplemented with propidium iodide was added to the cells. The DNA content of the cells was measured by flow cytometry using a 488 nm laser (BD FACSVerse with BD FACSuite software, version 1.0.6.). The cell cycle distribution was analysed with ModFit LT (Verity Software House, version 4.1.7).

### Colony formation

Cells were seeded at a density of 5000 cells/ml in 6-well plates and allowed to adhere overnight. Cells were then treated with compounds and incubated for 10 days. After the treatment, colonies were fixed with 70% ethanol, washed with PBS and stained with crystal violet. After 1 hour of incubation at RT, excess stain was removed by washing with PBS and distilled water, and the stained cell colonies were imaged by scanning. Colony formation was quantified by measuring the absorbance (570 nm, Infinite 200 Pro microplate reader, Tecan, Life Sciences) of crystal violet after solubilization with 1% SDS.

## Results

### 1. Anticancer effect of BTK inhibitors in breast cancer cell lines in vitro

The anticancer effect of the BTK inhibitors ibrutinib, acalabrutinib, tirabrutinib, evobrutinib and zanubrutinib was evaluated in a panel of ten breast cancer cell lines *in vitro*. The cell lines were categorized into two groups according to the reported expression status of the receptor: HER2-positive and HER2-negative [24, 25]. HER2 amplification and protein expression in the panel of cell lines were confirmed by FISH and western blot analysis, respectively (Supplementary Fig. 1).

**Fig. 1.**
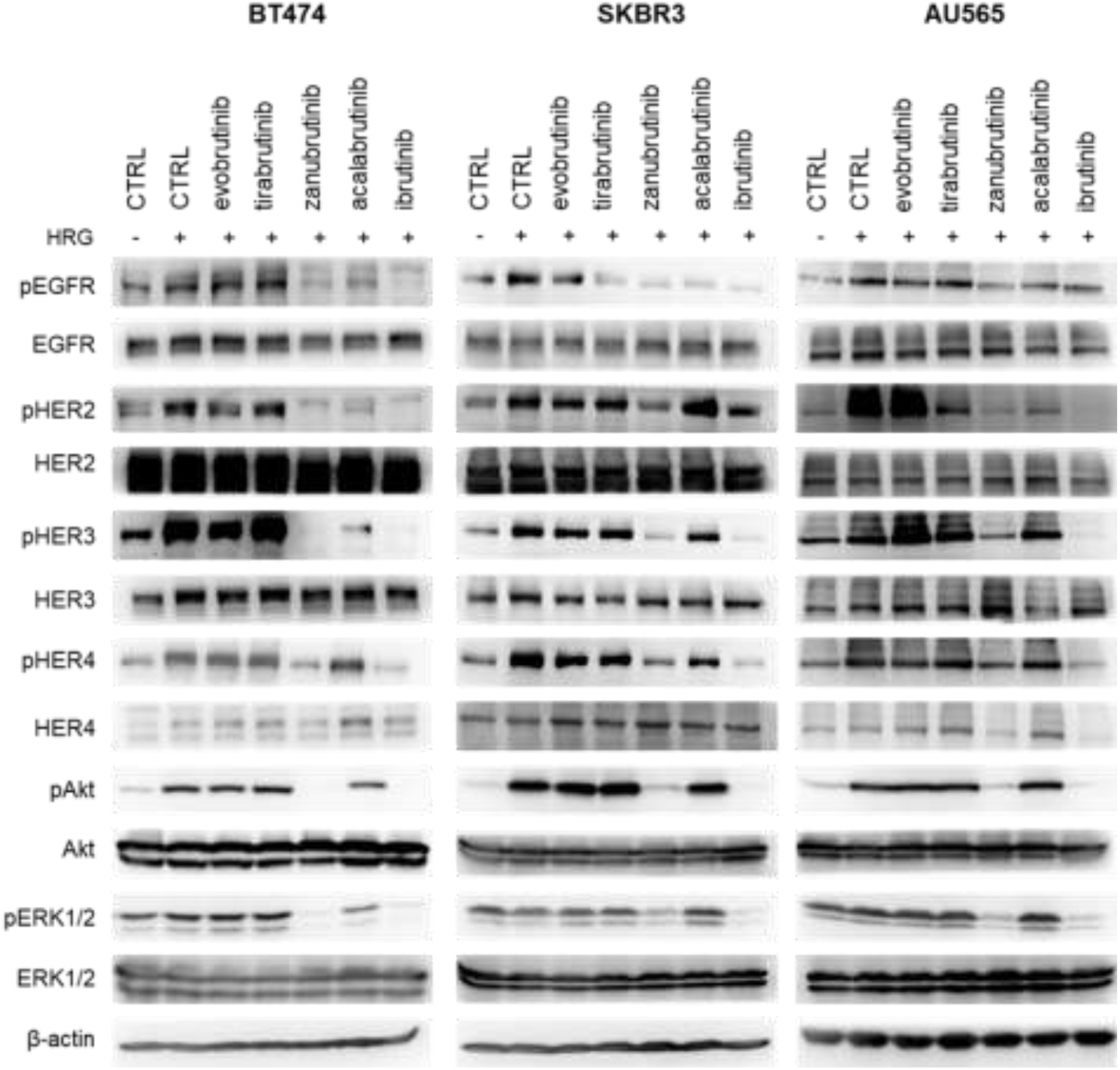
Effects of BTK inhibitors on signalling pathways in HER2-positive breast cancer cell lines. Compounds were used at a 10 μM concentration for 16 hours of treatment. Cells were stimulated by heregulin (HRG, 0.1 μg/mL) 30 min prior to harvesting. β-Actin served as a control for equal loading. Next, the dose-dependent effect of zanubrutinib was examined in BT474 and SKBR3 cells treated for 16 hours. Activation of ERBB receptors was impaired starting at a 1 μM concentration of the compound in BT474 cells, and the effect was dose related (Fig. 2).

The effect of the BTK inhibitors in the cell line panel was investigated, and GI50 values were determined after 72 hours of treatment using a resazurin assay to assess viability. Lapatinib was used as a positive control. The HER2-negative cell lines (MCF7, T47D, BT20, HCC1806) were insensitive to treatment with all compounds. In contrast, four HER2-positive cell lines were found to be sensitive to ibrutinib, zanubrutinib and acalabrutinib, with submicromolar GI50 values ranging from 0.09 μM to 0.88 μM for ibrutinib and zanubrutinib and single-digit micromolar GI50 values for acalabrutinib. The viability of the HER2-positive cell line JIMT-1 was not affected by the tested compounds, but these cells are known to be trastuzumab resistant [25]. Interestingly, the novel BTK inhibitor zanubrutinib was active at concentrations comparable to lapatinib. The activity of ibrutinib was approximately 1.5-8 times higher than that of lapatinib, which is in agreement with previously published data [19].

### 2. Zanubrutinib and acalabrutinib inhibit ERBB signalling in HER2-positive breast cancer cell lines

The effect of the tested BTK inhibitors on ERBB signalling in cancer cell lines stimulated by heregulin was analysed. Heregulin is the most broadly active ERBB ligand in HER2-positive breast cancer cells [26]. Heregulin is a potent activator, especially of HER3 and HER4, and possesses mitogenic activity in breast cancer cells [27, 28].

Specifically, the activating autophosphorylation of ERBB2 and subsequent phosphorylation of the downstream kinases Akt and ERK1/2 were analysed in cells treated with BTK inhibitors at a single concentration of 10 μM for 16 hours. According to previous reports, ibrutinib impaired the phosphorylation of ERBB receptors and the downstream kinases Akt and ERK1/2 [19, 29, 30]. Among the other compounds, zanubrutinib was similarly effective. Acalabrutinib and tirabrutinib showed slight inhibitory effects against EGFR and HER2 in some cell lines. Treatment with evobrutinib did not cause any changes in the phosphorylation of the evaluated proteins, probably due to its selectivity profile with minimum effects towards ERBB receptor kinases [31]. In the HER2-negative cell line MCF7, no changes in the phosphorylation of the ERBB downstream target (pERK1/2 T202/Y204) were detected (Supplementary Fig. 2).

**Fig. 2.**
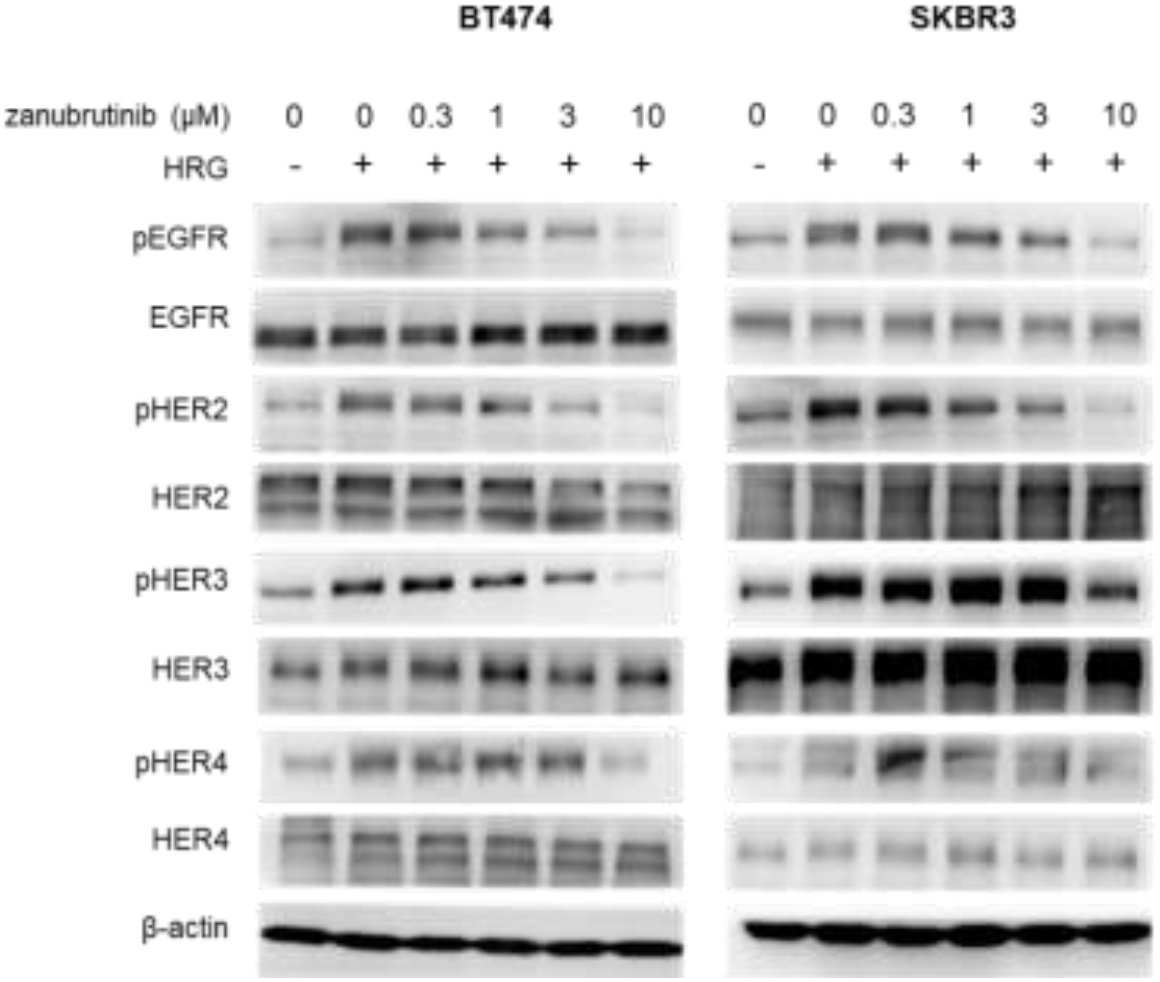
Dose-dependent effect of zanubrutinib in HER2-positive BT474 and SKBR3 cell lines. Cells were treated with increasing concentrations of zanubrutinib for 16 hours. Cells were stimulated by heregulin (HRG, 0.1 μg/mL) 30 min prior to harvesting. β-Actin served as a control for equal loading.

### 3. BTK inhibitors cause G1 arrest in cell lines overexpressing HER2

Cell cycle arrest at the G1-S transition induced by ibrutinib has been previously reported [19, 29]. As expected, zanubrutinib was found to have a similar effect on cell cycle progression in the HER2-positive cell lines BT474 and SKBR3. The compound increased the number of cells in G1 phase at higher concentrations than did ibrutinib (Fig. 3). The potent and specific EGFR/HER2 inhibitor lapatinib, used as a positive control, also resulted in accumulation of cells in G1 phase, although at much lower concentrations. In control experiments, no changes in the cell cycle were observed in HER2-negative MCF7 cells upon exposure to these three tested compounds (Supplementary Fig. 3).

**Fig. 3.**
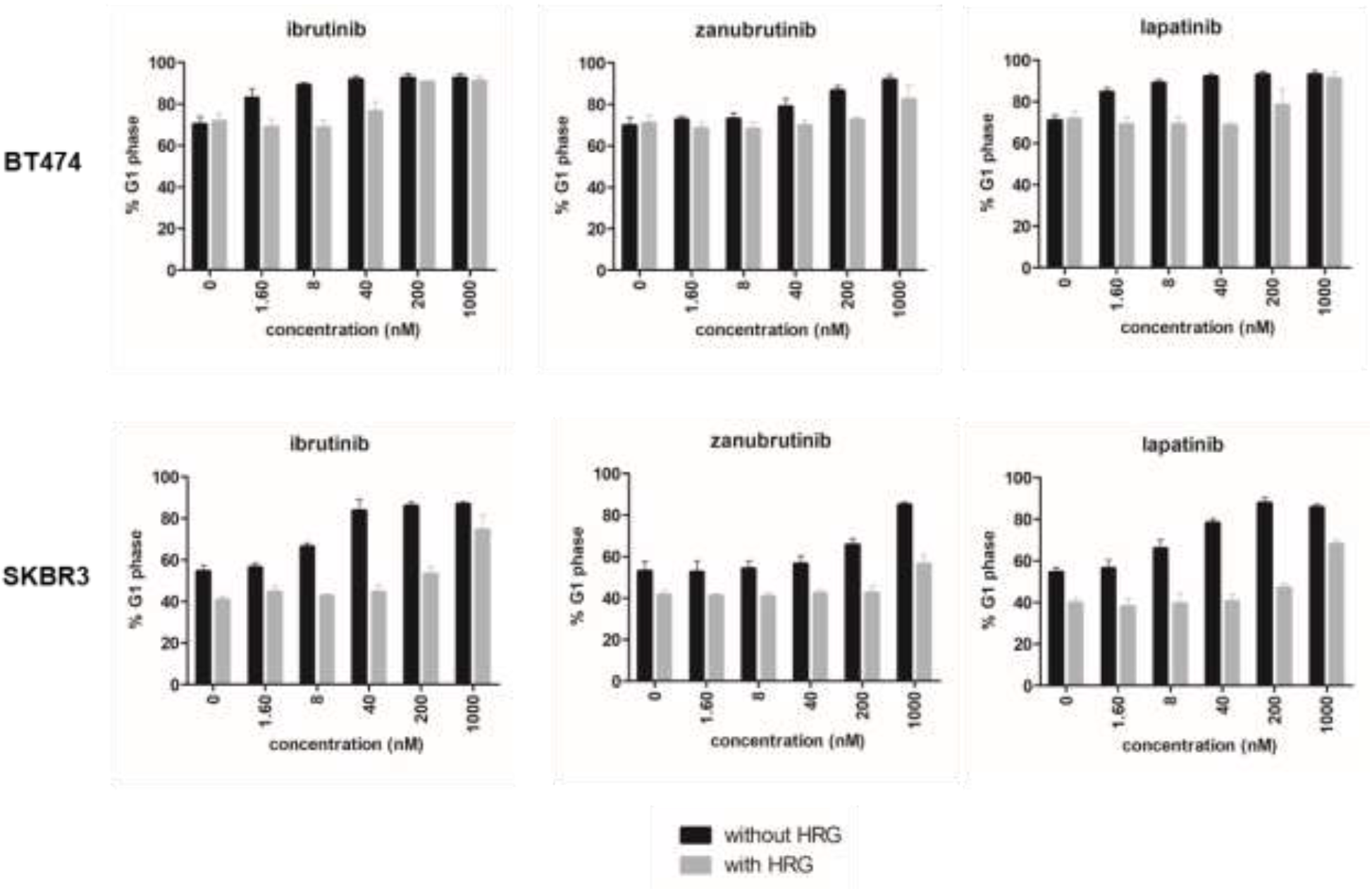
Heregulin rescues HER2-positive cells from G1 arrest induced by BTK inhibitors. Cells were treated for 24 hours with increasing concentrations of the selected compounds, with or without activation by heregulin (HRG, 0.1 μg/ml).

### 4. Heregulin impairs the effect of BTK inhibitors on the cell cycle distribution

We hypothesized that the addition of heregulin to cells exposed to BTK inhibitors would decrease the potency of these inhibitors and thus help to confirm that their mechanism of action is related to ERBB inhibition. We first analysed the effect of heregulin on the growth and viability of HER2-positive cells treated with ibrutinib and zanubrutinib for 72 hours. The results showed that cell growth and viability were restored when the treated cells were exposed simultaneously to heregulin (Table 2).

**Table 1.** Anticancer effects of BTK inhibitors in breast cancer cell lines

| | | GI<sub>50</sub> (μM)<sup>a</sup> | | | | | |
| --- | --- | --- | --- | --- | --- | --- | --- |
|  | cell line | ibrutinib | zanubrutinib | acalabrutinib | evobrutinib | tirabrutinib | lapatinib |
| HER2<sup>+</sup> | BT474 | 0.09 ± 0.01 | 0.81 ± 0.18 | 1.41 ± 0.18 | >5 | >5 | 0.33 ± 0.03 |
| | SKBR3 | 0.13 ± 0.02 | 0.84 ± 0.15 | 1.91 ± 0.16 | >5 | >5 | 0.47 ± 0.15 |
| | AU565 | 0.31 ± 0.14 | 0.73 ± 0.03 | 2.03 ± 0.49 | >5 | >5 | 0.76 ± 0.23 |
| | EFM192A | 0.10 ± 0.02 | 0.88 ± 0.10 | 1.41 ± 0.15 | >5 | >5 | 0.84 ± 0.20 |
|  | JIMT1<sup>b</sup> | >5 | >5 | >5 | >5 | >5 | >5 |
| HER2<sup>-</sup> | BT20 | >5 | >5 | >5 | >5 | >5 | >5 |
|  | HCC1806 | >5 | >5 | >5 | >5 | >5 | >5 |
|  | PMC42 | >5 | >5 | >5 | >5 | >5 | >5 |
|  | MCF7 | >5 | >5 | >5 | >5 | >5 | >5 |
|  | T47D | >5 | >5 | >5 | >5 | >5 | >5 |

<sup>a</sup>GI<sub>50</sub> values were obtained after 72 hours of treatment. <sup>b</sup> Trastuzumab insensitive cell line. GI<sub>50</sub> > 5 $\mu$ mol/L is considered inactive to ibrutinib based on previous reports [19]

**Table 2.** The cytotoxic effect of ibrutinib and zanubrutinib is dependent on heregulin

| Cell line | $GI_{50}$ ($\mu M$)<sup>a</sup> | | | |
| --- | --- | --- | --- | --- |
|  | ibrutinib | ibrutinib + HRG | zanubrutinib | zanubrutinib + HRG |
| BT474 | 0.09 ± 0.01 | >5 | 0.81 ± 0.18 | >5 |
| SKBR3 | 0.13 ± 0.02 | >5 | 0.84 ± 0.15 | >5 |

<sup>a</sup> Values were obtained after 72 hours of treatment

Next, we tested the effect of different concentrations of ibrutinib, zanubrutinib, and lapatinib on the cell cycle in HER2-positive cells stimulated with heregulin. Cotreatment with heregulin clearly rescued BT474 cells from G1 phase arrest at lower concentrations of the compounds (Fig. 3). The effect of the inhibitors was delayed under cotreatment with heregulin in comparison with treatment with each compound alone. The ability of heregulin to rescue HER2-positive breast cancer cells from the growth inhibition induced by ibrutinib and lapatinib is consistent with other findings [29].

### 5. Ibrutinib and zanubrutinib suppress colony formation and induce apoptosis in HER2-positive breast cancer cells

To further confirm the anticancer effect of zanubrutinib in the HER2-positive cell lines SKBR3 and BT474, we performed a long-term treatment experiment. SKBR3 cells showed a reduced ability to form colonies in the presence of zanubrutinib in a dose-dependent manner (Fig. 4A), although the effect was weaker than that of ibrutinib [28]. In agreement with the insensitivity of HER2-negative breast cancer cells to the studied BTK inhibitors (Table 1), the control cell line MCF7 also showed no decrease in colony formation after treatment with any of the compounds.

**Fig. 4.**
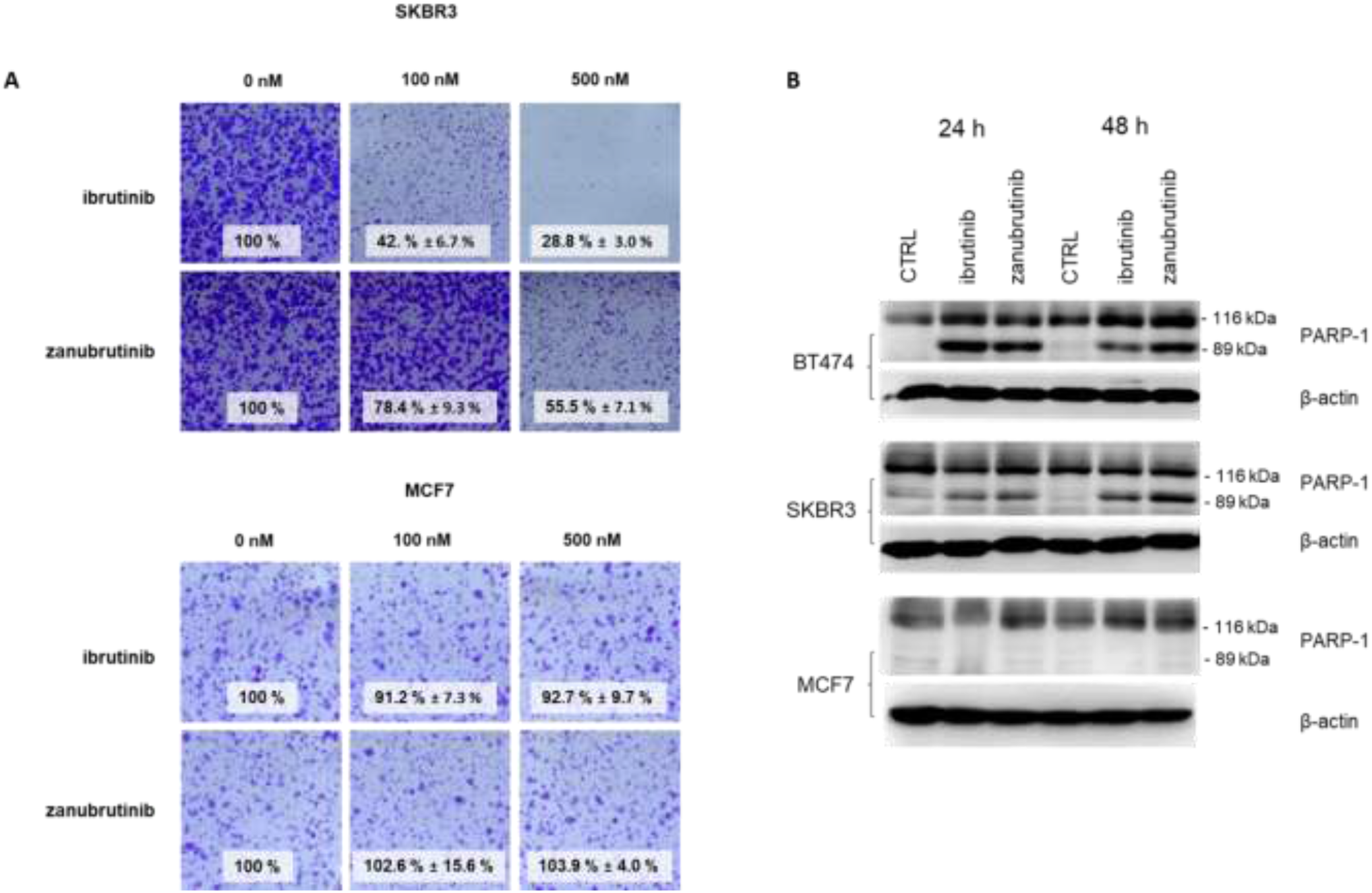
(A) Ibrutinib and zanubrutinib inhibit colony formation in HER2+ SKBR3 cells, while the proliferation rate of HER2-MCF7 cells remains unaffected. Colonies were stained with crystal violet after 10 days of treatment. The numbers indicate the percentages of colonies formed (calculated from the absorbance values). (B) Immunoblot analysis of lysates of cells treated with 1 μM zanubrutinib and ibrutinib. β-Actin served as a control for equal loading.

BTK inhibitors not only block the proliferation of HER2-positive cells but also directly induce their death. The level of cleaved PARP-1 was investigated in SKBR3, BT474 and MCF7 cells. Treatment with zanubrutinib and ibrutinib resulted in an increase in the cleaved PARP-1 fraction after 24 h in HER2-positive cell lines (Fig. 4B). In contrast, no change in the PARP-1 level was observed in HER2-negative MCF7 cells.

## Discussion

A current trend and attractive subject of research in drug development is the reuse of already known and approved drugs for new diseases, a strategy known as drug repurposing. Although modern approaches in novel drug development allow the design and dynamic development of potential pharmaceuticals, the search for suitable candidate compounds and their subsequent translation into therapies remains a relatively high hurdle. In the case of drug repurposing strategies, the application of established drugs for other indications brings with it some advantages. These include the already known safety profiles of the drugs, the acceleration of the approval process, and the consequent reduction in costs.

Ibrutinib and its follow-up BTK inhibitors are being investigated in a variety of clinical trials beyond their primary indications approved by the FDA. The inclusion of the drugs in preclinical tests utilizes both the primary target, BTK, and off-target kinases possessing a homologous Cys481 residue in the active site of the enzyme (e.g., EGFR, HER2, ITK) [17]. For ibrutinib and acalabrutinib, several clinical trials in solid tumours are ongoing. Zanubrutinib, the most recently approved BTK inhibitor, has not yet been evaluated for repurposing for solid tumours, although its biochemical and pharmacokinetic properties suggest possible activity towards HER2-overexpressing cancers. In our study, we therefore aimed to examine the potential repurposing of zanubrutinib and other selected BTK inhibitors in breast cancer cell line models.

Among the drugs tested in this study, ibrutinib and zanubrutinib demonstrated clear effects on cell models of HER2-positive breast carcinomas, as expected. The efficiency of ibrutinib in these cancers has already been proposed in previous publications [19, 29, 30], and we indeed found ibrutinib to be the most potent BTK inhibitor of ERBB signalling in breast cancer cell lines, once again confirming its anticancer effects. The potency of other tested BTK inhibitors against HER2-positive cell lines reflects the previously reported inhibition of individual ERBB receptors. Most interestingly among these other BTK inhibitors, zanubrutinib has been proven to inhibit the activity of EGFR and HER4 (86 and 86%, respectively) and of HER2 by 40% at 1 μM [22]. Our findings are in agreement with previously stated inhibitory potencies and reveal zanubrutinib as an effective inhibitor of proliferation and signalling in HER2-positive breast cancers.

We demonstrated that zanubrutinib effectively reduced the levels of phosphorylated forms of ERBB receptors in the HER2-overexpressing cell lines BT474, SKBR3, AU565 and EFM192A. Subsequently, downstream signalling of the Akt and ERK pathways was impaired, leading to reduced proliferation of the cells and proapoptotic effects. The effect of zanubrutinib was detectable when cells were treated with 3 μM zanubrutinib overnight (Fig. 2). The importance of the HER2 receptor in signalling and survival in HER2-positive breast cancer cell lines was supported by the strong antiproliferative effect of ibrutinib and zanubrutinib at submicromolar concentrations in these cell lines compared to cells with low expression of HER2, which were insensitive to BTK inhibitors.

In the presence of zanubrutinib, HER2-positive breast cancer cells exhibited G1 arrest. The deregulation of cell cycle progression in HER2-amplified cells reflects the detected inhibitory effect of BTK inhibitors on the phosphorylation of proteins in ERBB receptor-controlled pathways. G1 arrest was also observed in cells treated with ibrutinib and lapatinib, which were used as controls, suggesting that zanubrutinib acts via a similar mechanism [29].

In previous studies, the BTK-C transcript was detected in HER2-positive breast cancer cells [32]. Based on that finding, it was proposed that BTK-C signalling could be involved in the appearance of ligand-dependent lapatinib resistance in HER2-positive breast cancer cells and thus may be a potential therapeutic target in combination with HER2 in this subtype of breast carcinoma [28]. The molecular weight of BTK-C is 79.9 kDa [32], and its expression is detectable using a commercial antibody against BTK-A; however, in our study, we were not able to detect the expression of BTK in any of the tested breast cancer cell lines (Supplementary Fig. 1). Moreover, the study by Eifert et al. revealed BTK expression in a HER2-negative cell line and showed increased levels of apoptosis in BTK knockdown cells, whether HER2-positive or HER2-negative [31]. Thus, the presence of BTK in breast cancer cells may contribute to the effect of BTK inhibitors in HER2-positive cells. However, we assume that the effect of ibrutinib and zanubrutinib in this breast cancer subtype presumably stems from inhibition of ERBB signalling, as we did not observe any antiproliferative effects of the compounds in HER2-negative cells.

In conclusion, our findings support the potential indication of the BTK inhibitor zanubrutinib for use in HER2-amplified breast carcinomas. The anticancer effects of zanubrutinib have been described *in vitro*, and we suggest the compound as an attractive drug to be tested in further trials and *in vivo* models for the possibility of its repositioning outside of haemato-oncological diseases.

## Supplementary Information

The online version contains supplementary material available at …

## Acknowledgement

We thank Vladimira Koudelakova for performing FISH analysis.

## Author Contributions

All authors contributed to the study conception and design. Material preparation, data acquisition and analysis as well as preparing the first draft was performed by H.D., data analysis and writing of the manuscript was done by all authors. All authors read and approved the final manuscript

