## Supplementary material for "Anticancer effect of zanubrutinib in HER2-positive breast cancer cell lines"

| | |
|---|---|
| Supplementary table 1: | Percent inhibition of ERBB kinases by BTK inhibitors (1 μM) |
| Supplementary figure 1: | HER2 and BTK status verification of chosen breast cancer cell lines either via immunoblotting (A) or FISH (B) |
| Supplementary figure 2: | Effects of BTK inhibitors on ERBB downstream target in HER2 negative breast cancer cell line MCF7 |
| Supplementary figure 3: | BTK inhibitors show no significant impact on cell cycle progression in HER2 negative cell line MCF7 |

**Supplementary table 1**. Percent inhibition of ERBB kinases by BTK inhibitors (1 µM). Data extracted from published literature (Guo, Y. et al. (2019) & Crawford, J. J. et al., (2018))

| **inhibitor** | **EGFR** | **HER2** | **HER4** |
|---|---|---|---|
| ibrutinib[23] | 89 | 91 | 94 |
| acalabrutinib[23] | -1 | 36 | 93 |
| zanubrutinib[22] | 86 | 40 | 96 |
| tirabrutinib[23] | -2 | 7 | 28 |
| evobrutinib[23] | 1 | 1 | 57 |

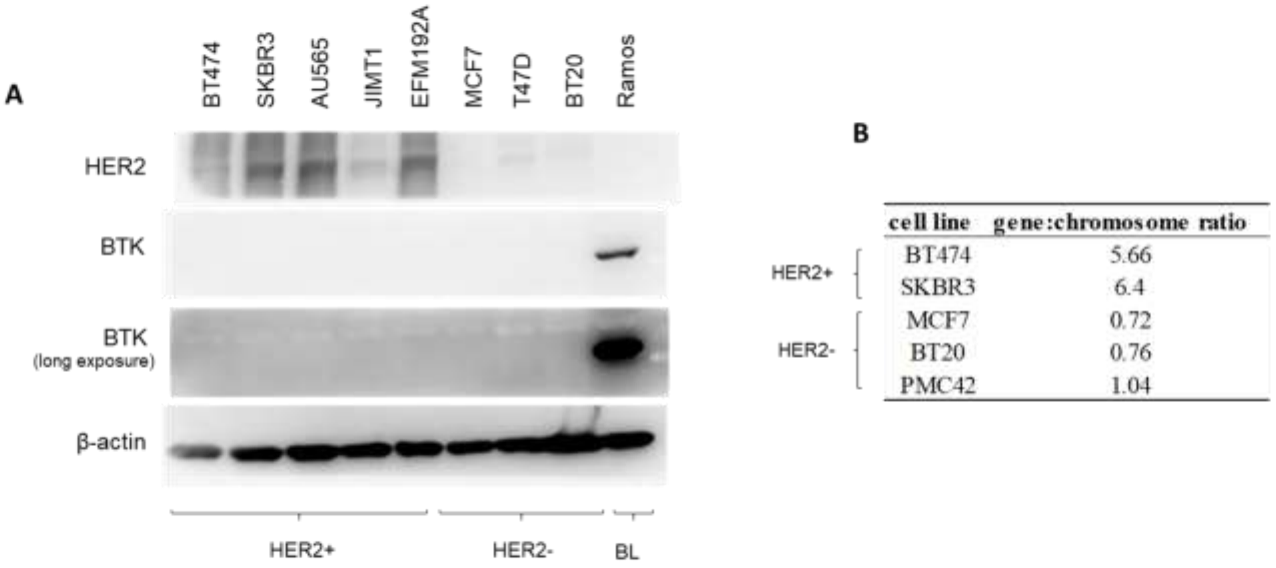


**Supplementary fig. 1** HER2 and BTK status verification of chosen breast cancer cell lines either via immunoblotting (A) or FISH (B). β-Actin served as control of equal loading. BL –Burkitt's lymphoma.

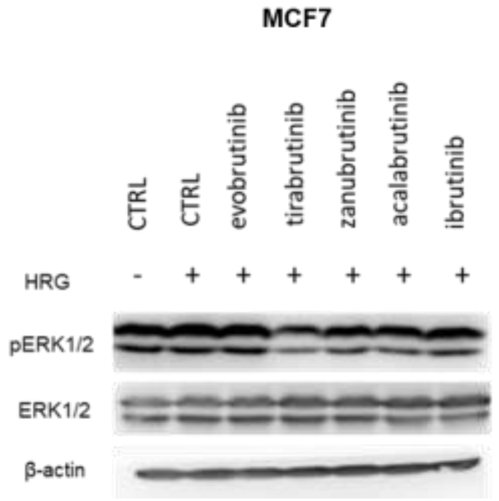


**Supplementary fig. 2** The effects of BTK inhibitors on ERBB downstream target in HER2 negative breast cancer cell line MCF7. Compounds were used in 10 μM concentration for 16 hour treatment. Cells were stimulated by heregulin (HRG, 0.1 μg/ml) 30 min prior to harvesting. β-Actin served as control of equal loading.

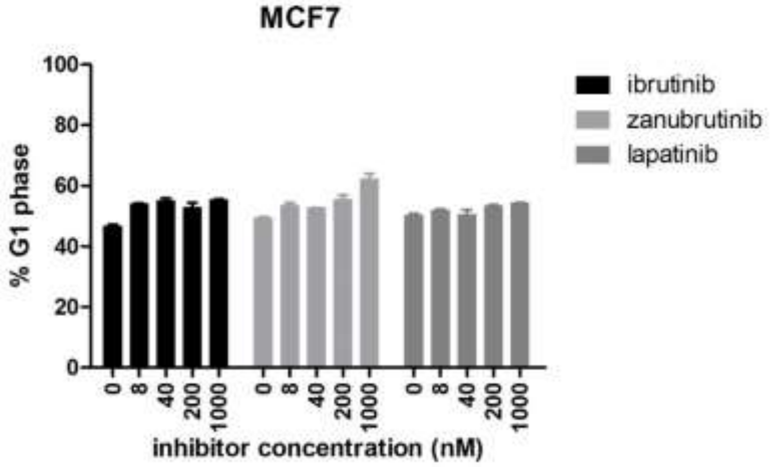


**Supplementary fig 3** BTK inhibitors show no significant impact on cell cycle progression in HER2 negative cell line MCF7. Cells were treated with indicated concentrations of selected compounds for 24 hours.
